# Nutritional value of extracted soybean, full fat soya and maize gluten meal proteins in comparison to a fish meal based reference diet for fingerling tilapia (*Oreochromis mossambicus*)

**DOI:** 10.1101/2023.11.12.566624

**Authors:** M. E. Bell, S. J. Davies

**Affiliations:** School of Biological and Marine Sciences (Faculty of Science and Engineering), Drake Circus, University of Plymouth, Plymouth, UK, PL4 8AA; Aquaculture and Nutrition Research Unit (ANRU), Carna Research Station, Ryan Institute and School of Natural Sciences, University Galway, Connemara, Co. Galway, H91 V8Y1, Ireland

**Keywords:** tilapia, soybean, maize gluten, fish meal comparison, growth, feed efficiency

## Abstract

A two-month investigation evaluated potential selected plant protein sources compared to fish meal protein in scientifically formulated diets for fingerling tilapia (*Oreochromis mossambicus*). Six nutritionally balanced diets: one reference (control) diet (fish-meal-based diet), three selected plant proteins (solvent extracted soybean, full fat soybean, and maize gluten) were substituted in the LT (Low Temperature dried) fish-meal-based control diet protein at 50% respectively, and the remaining diets were replaced with 75% inclusion of solvent extracted soybean and maize gluten meals. Results showed that SESB50, FFSB50 and MG50 fed tilapia did not appreciably differ in their growthperformance (final mean weights of 12.07, 10.75 and 9.18% respectively) with a final mean weight of 14.95g being significantly superior for the fish meal control (CO) group. Substitution of fish meal diet with solvent-extracted soybean and maize gluten at levels of 75% gave the poorest growth performance and Feed Efficiency (FE) (7.01g final weight, 51% FE). However, the MG50 diet also showed some inferior feed utilisation performance. However, tilapia fed on a full fat soybean diet (FFSB50) did not vary significantly from the control (CON) diet in terms of Apparent Net Protein Utilisation (ANPU) at 34.20 and 37.53% respectively. Whilst this preliminary study concluded that limitations on the use of the selected plant proteins in diets for tilapia are apparent, several approaches that could result in the future improvement of the nutritional value of soybean and maize gluten products for use in fish diets are also stated.

**Highlights:** A scientifically formulated reference diet using a Low Temperature dried (LT) fish meal produced superior growth performance for fingerling tilapia compared to selected oilseed (soybean) and grain (maize gluten) derived protein concentrates.

Soybean and full fat soybean meals were effective in contributing to 50% inclusion in experimental diets, but 75% of soybean and maize gluten meal was not able to support similar growth of tilapia over eight weeks.

Feed utilisation parameters such as Protein Efficiency Ratio (PER), Apparent Net Protein Utilisation (ANPU), Net Energy Utilisation (NEU) and Feed Efficiency (FE) gave supporting evidence and conclusions for optimal threshold inclusions of the test plant ingredients for tilapia.

## Introduction

Aquaculture is considered a growing and significant part of the global agribusiness and seafood industry. Fish farming operations produced more than 18 million tons in 2000 to 82 million tons in 2020, making it the fastest-growing food production sector in the world (FAO, 2020). Aquaculture is an important development in most of Southeast Asia, particularly within China. Freshwater production accounts for one-third of the total aquaculture in Southeast Asia (Kayansamruaj *et al*., 2020). This corresponds to the increased consumer demands from a growing population for seafood products. Asia remains the leading producer within the aquaculture domain globally, generating 92% of live-weight volume in 2017 (Naylor *et al*., 2021). From this, 75% of output is derived from freshwater aquaculture, evidencing the global trends and scientific breakthroughs using sustainable aquifers and feed technologies (Naylor *et al*., 2021). In developing countries, their objective is to produce high-value animal protein which cannot be supplied by traditional culture in sufficient quantity to support the ever-growing population, now over 8 billion as of 2022, and compensate for the desertification of the land (Barnabe, 1994; UN, 2022). Common species that are being cultured for commercial purposes are seabass, tilapia, milkfish, snakehead, catfish, eels, and carp. Such species are a valuable addition to their capture fishing methods and the technology for fish culture is now an important part of the economy of these regions. Of the many fish under consideration, tilapia is the most widespread and is a major tropical fish for intensive production.

Several investigations have addressed the effectiveness of replacing fish meal with some other protein source, particularly plant proteins. It has generall y been found that most alternative protein sources are able to replace fish meal to varying extents (De. Silva, 1998). Several factors affect the proportion of fish meals that can be replaced in balanced feeds. This inclusion depends upon the nature of the protein source and essential amino acid profiles. Fish meal replacements have numerous shortcomings, often including low protein content and amino acid levels that do not meet requirements, low energy content per unit weight, poorer digestibility, inferior palatability to fish, and consumer acceptability (Sadiku and Jauncey, 1995). Gule and Geremew (2022) extensively reviewed the use of novel and traditional ingredients for finfish aquaculture. Likewise, El-Sayed (1999) and Shiau (2002) have reviewed tilapia nutrition, feed requirements, and dietary specifications.

Aquafeeds still utilises significant fish meal and fish oil sourced from wild-captured forage fish to meet the nutrient specifications of farmed fish. However, the increasing use of forage fish is unsustainable and because an additional 37.4 million tons of aquafeeds will be required by 2025, competing protein sources are being employed to meet this emerging ‘protein gap’. Fish meal is now used strategically in most aquafeeds.

Beyond plant-based ingredients, various novel byproducts like PAPS (processed animal proteins) and insect meals, algae, and Single-cell proteins (SCPs) have much potential to supply the protein aquafeeds required over the next 10–20 years. However, for mainstream oilseed meal and grain-based sources (soya and cereal proteins), we require more accurate threshold constraints to better formulate practical diets using such approaches as linear least-cost formulation (LLF). Therefore, this study aimed to evaluate the growth response of tilapia, to which a substantial proportion of a Low Temperature dried (LT) fish meal was replaced with selected plant proteins. The experimental strategy for this investigation was to test plant ingredients against the fish meal as a high-quality Biological Value (BV) standard to obtain better resolution values for precision diets for tilapia. The plant proteins that were used in this experiment were solvent-extracted soybean meal, full fat soybean, and maize gluten respectively. These traditional protein sources were tested at various inclusion rates compatible with providing 38% protein and 12% lipid in a balanced diet at the expense of a low temperature fish meal as a high Biological Value (BV) reference protein.

## Materials and Methods

### Experimental diets

Six experimental diets were formulated to contain a variable proportion of plant protein to replace fish meal partially at fixed levels. The experimental diets were formulated by using a linear least-cost software application FeedSoft ™. All diets were made iso-caloric and iso-nitrogenous and were adjusted at appropriate levels and to contain 38% crude protein and 12% lipid. The proximate feed formulation and chemical composition of the experimental diets are depicted in Table 1. The proximate chemical composition of the used protein sources is presented in Table 3 and these values were used in the formulation software. A control diet (CON) was based on LT (Low Temperature Dried) fish meal as the main source of dietary protein, and test ingredients were solvent-extracted soybean meal (SESB), full fat soybean (FFSB), and maize gluten (MG) respectively.

**Table 1.**
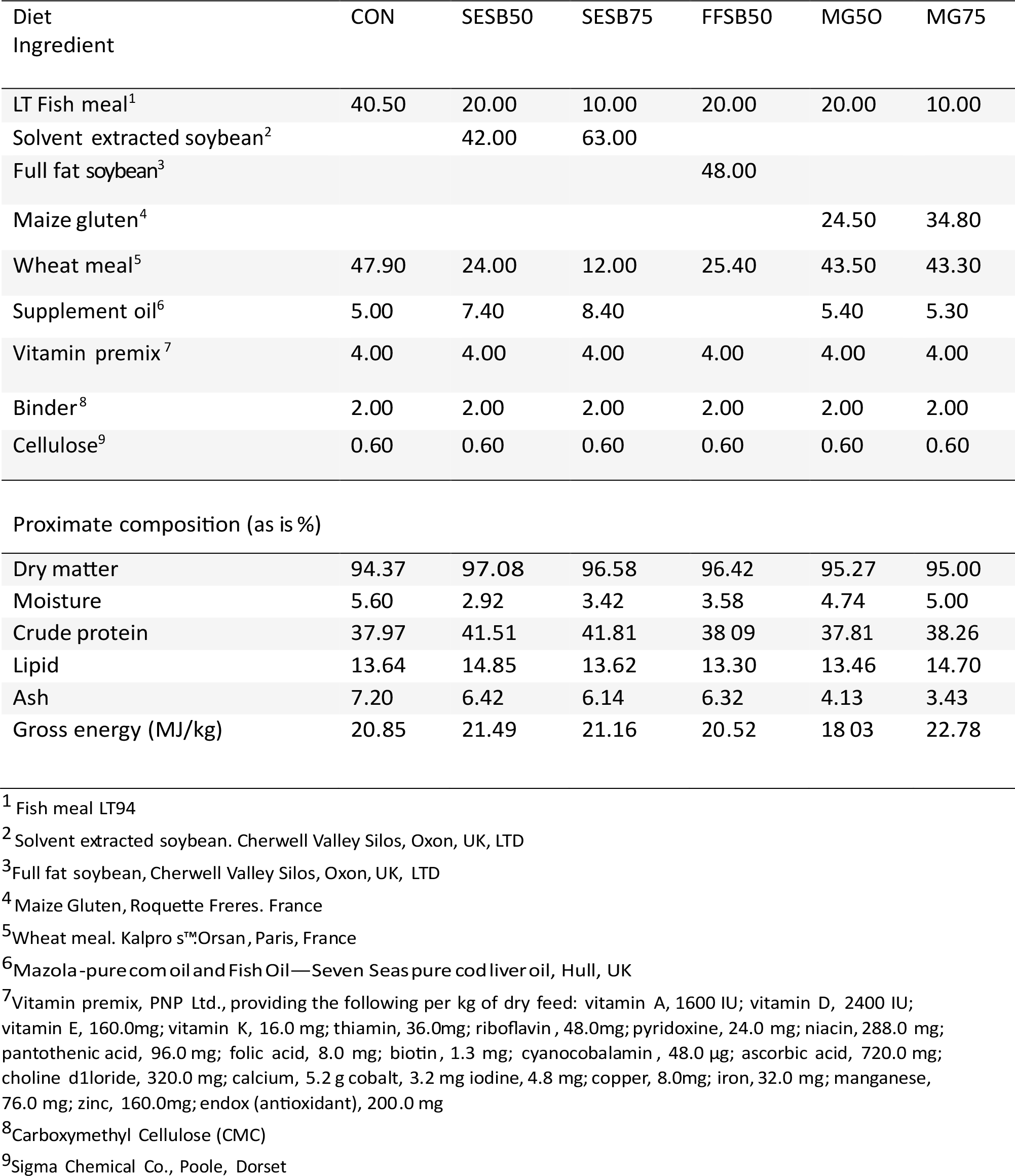
Feed formulation (as %) and proximate analysis results of experimental diets (as %)

| Diet<br>Ingredient | CON | SESB50 | SESB75 | FFSB50 | MG50 | MG75 |
| --- | --- | --- | --- | --- | --- | --- |
| LT Fish meal<sup>1</sup> | 40.50 | 20.00 | 10.00 | 20.00 | 20.00 | 10.00 |
| Solvent extracted soybean<sup>2</sup> |  | 42.00 | 63.00 |  |  |  |
| Full fat soybean<sup>3</sup> |  |  |  | 48.00 |  |  |
| Maize gluten<sup>4</sup> |  |  |  |  | 24.50 | 34.80 |
| Wheat meal<sup>5</sup> | 47.90 | 24.00 | 12.00 | 25.40 | 43.50 | 43.30 |
| Supplement oil<sup>6</sup> | 5.00 | 7.40 | 8.40 |  | 5.40 | 5.30 |
| Vitamin premix<sup>7</sup> | 4.00 | 4.00 | 4.00 | 4.00 | 4.00 | 4.00 |
| Binder<sup>8</sup> | 2.00 | 2.00 | 2.00 | 2.00 | 2.00 | 2.00 |
| Cellulose<sup>9</sup> | 0.60 | 0.60 | 0.60 | 0.60 | 0.60 | 0.60 |
| Proximate composition (as is %) |  |  |  |  |  |  |
| Dry matter | 94.37 | 97.08 | 96.58 | 96.42 | 95.27 | 95.00 |
| Moisture | 5.60 | 2.92 | 3.42 | 3.58 | 4.74 | 5.00 |
| Crude protein | 37.97 | 41.51 | 41.81 | 38.09 | 37.81 | 38.26 |
| Lipid | 13.64 | 14.85 | 13.62 | 13.30 | 13.46 | 14.70 |
| Ash | 7.20 | 6.42 | 6.14 | 6.32 | 4.13 | 3.43 |
| Gross energy (MJ/kg) | 20.85 | 21.49 | 21.16 | 20.52 | 18.03 | 22.78 |

124 <sup>1</sup> Fish meal LT94125 <sup>2</sup> Solvent extracted soybean. Cherwell Valley Silos, Oxon, UK, LTD126 <sup>3</sup> Full fat soybean, Cherwell Valley Silos, Oxon, UK, LTD127 <sup>4</sup> Maize Gluten, Roquette Freres. France128 <sup>5</sup> Wheat meal. Kalpro s™ Orsan, Paris, France129 <sup>6</sup> Mazola-pure com oil and Fish Oil—Seven Seas pure cod liver oil, Hull, UK
<sup>7</sup> Vitamin premix, PNP Ltd., providing the following per kg of dry feed: vitamin A, 1600 IU; vitamin D, 2400 IU; vitamin E, 160.0mg; vitamin K, 16.0 mg; thiamin, 36.0mg; riboflavin, 48.0mg; pyridoxine, 24.0 mg; niacin, 288.0 mg; pantothenic acid, 96.0 mg; folic acid, 8.0 mg; biotin, 1.3 mg; cyanocobalamin, 48.0 µg; ascorbic acid, 720.0 mg; choline d1loride, 320.0 mg; calcium, 5.2 g cobalt, 3.2 mg iodine, 4.8 mg; copper, 8.0mg; iron, 32.0 mg; manganese, 76.0 mg; zinc, 160.0mg; endox (antioxidant), 200.0 mg
<sup>8</sup> Carboxymethyl Cellulose (CMC)136 <sup>9</sup> Sigma Chemical Co., Poole, Dorset

Three diets were formulated in which 50% of the fish meal diet was replaced by selected plant proteins, i.e., SESB50, FFSB50 and MG50. Whereas for the remaining diets, 75% of the substitution of fish meal diet with solvent-extracted soybean and maize gluten (SESB75 and MG75) was done. Wheat meal was also included as the main carbohydrate energy filler source for balance.

The diets were processed by blending the dry ingredients into a homogenous mixture with a Hobart A 120 industrial food processor. The required supplementary oil for each diet was added gradually and after a few minutes of mixing, 320ml of water was added to prime the binder. Once a homogenous dough mixture was obtained, the diets were extruded through a mincer into ‘spaghetti-like’ strands and broken into smaller pellets. Pellets were dried by convection air in an oven at 40°C. for 12 hrs. After cooling, the diets were packed in sealed airtight containers, and stored at −30°C until needed. Before feeding, the diets were further broken into smaller pellets (1-2mm diameter) to suit the gape size of the fish.

### Experimental systems and maintenance

The feeding trial was conducted in a RAS-controlled facility using 650 *Oreochromis mossambicus* obtained from FishGen Ltd (Wales UK), of average mean live weight 0.95g. Initially, the fry was fed on a sex-reversal diet (BioMar Incio + Trout Fry Diet) treated with 30ppm 17 α-methyl testosterone) for four weeks to ensure an almost complete male phenotype brood. This was done to avoid differences in growth rate between males and females at maturation, which occurs early in this species. Fry were acclimatized to the tank environment one week prior to the start of the experiment. Eighteen 24-litre self-cleaning fibreglass tanks were used with the continuous freshwater flow (2.4 l/min) through a mechanical and biological filter. The water in the main reservoir was maintained at approximately 26± 1°C by a thermostatically controlled immersion heater and vigorously heated. pH, NH_3_ NO_2_ and NO_3_, were monitored and remained at acceptable levels throughout the experimental period.

### Feeding regime

Initially, fish were fed for one week to acclimate them to the diet and the system and free their gastrointestinal tract from the pre-experimental diet. At the end of the acclimation period, the fish were weighed and subsequently started on experimental diets. Each of the 18 tanks was randomly assigned to respective dietary treatment (in triplicate groups). Thirty-six fish were graded and transferred to each tank with an average weight of 3.15g. A sub-sample group of 20 fish were euthanized using a lethal MS222 (Tricaine Methanesulfonate) solution and kept frozen (−18°C) to determine initial carcass composition.

Fish were fed twice daily, six days a week, at a feeding rate of 3% wet body weight per day. Fish were weighed on the seventh day. The feeding rate was increased to 4% from week five onwards as fish acclimated to a higher feeding plane. However, the feeding rates were adjusted to account for each diet’s moisture content and appropriate to changing biomass on a weekly basis (Table 3).

At the end of the experiment, the final weight of the fish was measured. Following a 24-hour starvation period, three groups of fish (n=3), where fifteen fish were collected from each treatment, were randomly selected from each experimental tank to analyse the carcass composition.

### Analytical procedures

Proximate analyses of diets (Table 1), ingredients (Table 2) and carcasses were made following the usual procedures (AOAC, 2023). Specific growth rate (SGR), the rate of growth of an animal, is a sensitive index of protein quality under controlled conditions being proportional to the supply of essential amino acids. Daily SGR can be calculated by using the formula:

**Table 2.** Nutrient content of experimental ingredients. Fish meal, Wheat meal, Solvent-extracted soybean, Full fat soybean, and Maize Gluten.

| Ingredients<br>Nutrient<br>content | Fish<br>meal | Wheat meal | Solvent extracted<br>soybean | Full fat<br>soybean | Maize<br>Gluten |
| --- | --- | --- | --- | --- | --- |
| Moisture | 9.81 | 12.23 | 12.56 | 12.34 | 7.84 |
| Dry matter | 90.19 | 87.77 | 87.44 | 87.66 | 92.16 |
| Protein | 73.24 | 15.00 | 43.05 | 37.62 | 70.82 |
| Lipid | 13.31 | 2.16 | 3.15 | 19.00 | 11.29 |
| Ash | 12.88 | 3.73 | 6.03 | 4.87 | 1.58 |
| Energy<br>(MJ/kg) | 15.50 | 17.28 | 19.50 | 22.04 | 26.33 |

**Table 3.** Feeding rates for experimental diets on a dry matter basis and corrected for the moisture contents of the diets (% body wt./day) for phase I (1-4 weeks); phase II (5-8 weeks)

| Week 1 - 4 |  |  |  |  |  |  |  |
| --- | --- | --- | --- | --- | --- | --- | --- |
| Feeding rate (%) | Diet | CON | SESB50 | SESB75 | FFSB50 | MG50 | MG75 |
| Dry matter |  | 3.00 | 3.00 | 3.00 | 3.00 | 3.00 | 3.00 |
| Moisture content (%) |  | 5.63 | 2.92 | 3.42 | 3.58 | 4.74 | 5.00 |
| As moist diet<sup>1</sup> |  | 3.18 | 3.09 | 3.11 | 3.11 | 3.15 | 3.16 |
| Week 5 - 8 |  |  |  |  |  |  |  |
| Feeding rate (%) | Diet | CON | SESB50 | SESB75 | FFSB50 | MG50 | MG75 |
| Dry matter |  | 4.00 | 4.00 | 4.00 | 4.00 | 4.00 | 4.00 |
| Moisture content (%) |  | 5.63 | 2.92 | 3.42 | 3.58 | 4.74 | 5.00 |
| As moist diet<sup>1</sup> |  | 4.24 | 4.12 | 4.14 | 4.15 | 4.20 | 4.21 |

<sup>1</sup>[feeding rate/ (100-moisture content) x 100]

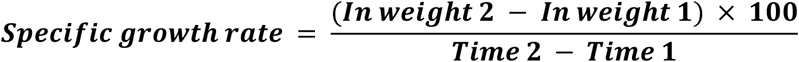

### Statistical analysis

The effect of diet was tested using analysis of variance. The results for initial and final fish weights were subjected to statistical analysis by one-way analysis of variance.Where significant differences at the 5% probability level were found between dietary treatments, the means were compared using Duncan’s Multiple Range Test. The data for carcass composition were based on pooled samples from each tank. From the mean initial and final weights and the values obtained for initial and final carcass protein content, Specific Growth Rate (SGR) and Apparent Net Protein Utilisation (NPU) for each dietary treatment could be calculated. Standard errors of means (± SE) were calculated to identify the range of means.

## Results

### Growth Performance

The growth response and feed utilisation data for tilapia fed the six experimental diets are displayed (Table 4). There was a significant difference between the final average body weight amongst the fish fed on the experimental diets. Fish fed on fish-meal-based control diets attained a 5-fold increase in final average body weight, which was 14.95g. However, the lowest value was observed for the fish fed the 75% inclusion level of maize gluten in the diet, replacing the fish meal component. Those fish had only approximately doubled their weights after eight weeks. The SESB50, FFSB50 and MG50 diets did not show any significant difference (P > 0.05) in the growth performance of fish respectively. Both SESB75 and MG50 diets did not show any appreciable difference in the final average body weight of the fish fed on those diets. These trends were seen also in the percentage weight gain figures, which decreased as the level of plant protein inclusion increased. The control diet supported the highest weight gain of 355.28%, while MG75% produced a weight gain of 126. 13%. SGR data further supported this trend, with SGR dropping from 2.72% per day for the control diet-fed fish to 1.46% per day for the fish fed MG75% diet. Fish-fed the 50% level inclusion of plant proteins performed better than those on the 75% level of plant proteins (Table 4). With equal quantities of protein occurring from plant and animal sources, a significant reduction in growth rate comparable with the fish fed on fish-meal-based control diets was observed. However, from week six onwards, fish fed on SESB75, MG50 and MG75 diets showed a depression growth response. The difference in growth rate manifested from week six onwards. At the end of the experimental weeks, fish fed on fishmeal-based diet was superior to the other experimental fish. No fish mortality occurred during the whole experimental period in any of the dietary treatments.

**Table 4.** Growth performance, feed utilisation efficiency of feed experimental diets. For 8-weeks.

| Diets | CON | SESB50 | SESB75 | FFSB50 | MG50 | MG75 | ±SEM<sup>1</sup> |
| --- | --- | --- | --- | --- | --- | --- | --- |
| **Parameters** |  |  |  |  |  |  |  |
| Initial mean weight (g) | 3.28 | 3.18 | 3.25 | 3.08 | 3.12 | 3.1 | 0.03 |
| Final mean weight (g) | 14.95<sup>a</sup> | 12.07<sup>b</sup> | 8.99<sup>bc</sup> | 10.75<sup>b</sup> | 9.18<sup>b</sup> | 7.01<sup>d</sup> | 1.13 |
| Specific growth rate (SGR %/day) | 2.71<sup>a</sup> | 2.38<sup>b</sup> | 1.82<sup>bc</sup> | 2.23<sup>b</sup> | 1.93<sup>c</sup> | 1.46<sup>c</sup> | 0.18 |
| Weight gain (%) | 355.18<sup>a</sup> | 279.56<sup>b</sup> | 176.62<sup>c</sup> | 249.03<sup>b</sup> | 194.23<sup>c</sup> | 126.13<sup>d</sup> | 33.37 |
| Feed efficiency (%) | 101.00<sup>a</sup> | 83.00<sup>b</sup> | 60.00<sup>c</sup> | 77.00<sup>bc</sup> | 69.00<sup>bc</sup> | 51.00<sup>d</sup> | 7.23 |
| Protein efficiency ratio (PER)* | 2.68<sup>a</sup> | 2.00<sup>b</sup> | 1.44<sup>d</sup> | 2.03<sup>b</sup> | 1.81<sup>c</sup> | 1.34<sup>d</sup> | 0.20 |
| Daily feed intake (g) | 8.69<sup>a</sup> | 8.03<sup>a</sup> | 7.17<sup>b</sup> | 7.46<sup>b</sup> | 6.64<sup>c</sup> | 5.42<sup>d</sup> | 0.46 |
| Mean voluntary feed intake (%) | 3.15 | 3.2 | 3.27 | 3.15 | 3.17 | 3.04 | 0.03 |
| Apparent net protein utilisation | 37.53<sup>a</sup> | 31.85<sup>b</sup> | 21.25<sup>d</sup> | 34.20<sup>a</sup> | 26.33<sup>c</sup> | 21.55<sup>d</sup> | 2.77 |
| Net energy utilisation (%) | 89.70<sup>a</sup> | 81.11<sup>b</sup> | 49.57<sup>e</sup> | 73.81<sup>c</sup> | 82.42<sup>b</sup> | 65.41<sup>d</sup> | 5.88 |

Note: Figures in each row having the same superscript are not significantly different (P < 0.05).
<sup>1</sup>Pooled standard error NS, not significant. \*PER= live weight gain (g)/Protein intake (g)

### Growth performance and feed efficiency

Feed intake and feed utilisation parameters are displayed in Table 4. Fish-fed SESB50 and FFSB50 diets did not differ (p < 0.05) significantly in final mean weight from each other (12.07-10.75g) but lower than the control group (CON) at 14.95g. Tilapia successfully converted 101% (FE) of their food to gain maximum weight to attain a final mean weight of 14.9g, but this reduced to just 51% for tilapia-fed MG75. Feed efficiency values all reflected the Specific Growth Rate (SGR) values (2.71-1.46% per day), with lower FE recorded for high replacement of fish-meal-based control diet with plant proteins. Tilapia fed maize gluten meal diets performed the worse with MG75 resulting in fish with a final weight of 7.01g after eight weeks with a percentage 126% weight gain compared to 355.18% for the control diet group.

A superior Protein Efficiency Ratio (PER) was observed for the diet containing the fish-meal-based (CON) diet (2.68), whereas the other diets did show significant (P > 0.05) differences concerning protein efficiency ratio for higher plant ingredients (2.00-1.34) (Table 4).

Tilapia fed on the fish meal-based control diet showed the best Apparent Net Protein Utilisation (ANPU) value but did not differ (P > 0.05) from the full fat soybean group (37.53% and 34.20% respectively). Feed consumption appeared to be significantly reduced as calorie density increased in diets at the given protein level and significantly affected growth response. Net Energy Utilisation (NEU) values also varied amongst the tilapia groups. Fish fed on the fish-meal-based control diet obtained the highest NEU (89.70%) and the lowest value (49.57%) was recorded for the SESB75 group (Table 4). NEU values reflected the ANPU data following overall nutrient assimilation.

### Carcass composition

The initial and final carcass composition of the tilapia fed on the experimental diets is presented in Table 5 after 8 weeks of feeding. The final carcass composition showed significant differences in their nutrient profiles. Tilapia fed on maize gluten MG50 and MG75 diets attained an appreciably high lipid content of 11.07% and 13.62%, respectively, compared to the fish meal control (CON) tilapia with a carcass level 8.76%. Fish fed on the fish-meal-based control diets, SESB75, MG50 and MG75 diets, showed no significant protein content variations (13.95-15.90%).

**Table 5.**
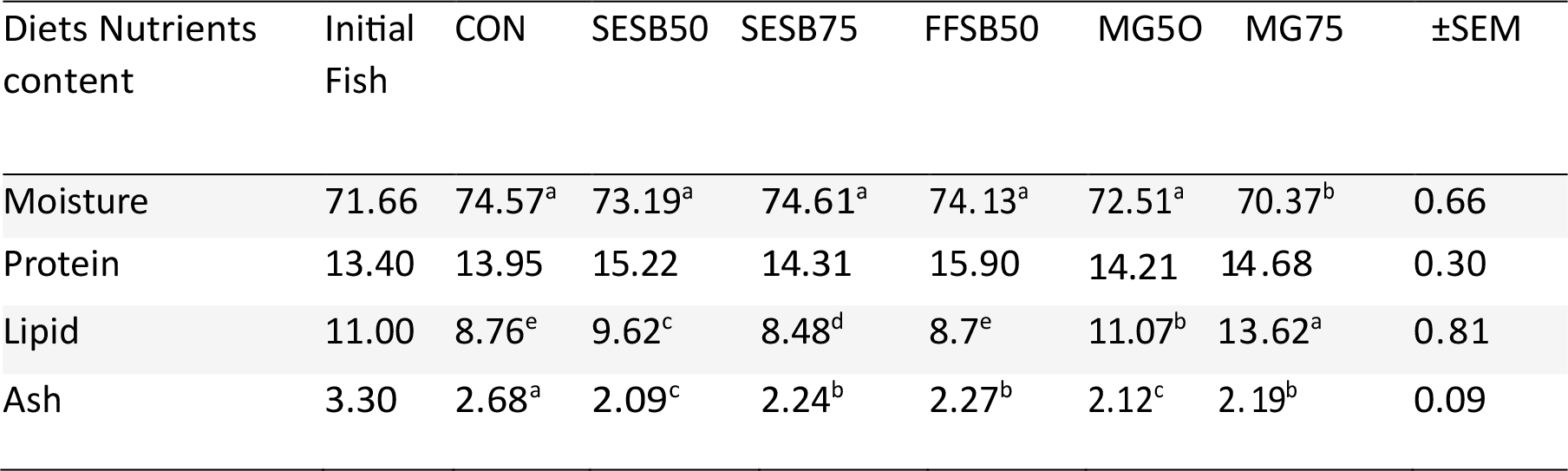
Carcass composition of *Oreochromis mossambicus* fed the experimental diets (as wet basis in percentage).

| Diets Nutrients content | Initial Fish | CON | SESB50 | SESB75 | FFSB50 | MG50 | MG75 | ±SEM |
| --- | --- | --- | --- | --- | --- | --- | --- | --- |
| Moisture | 71.66 | 74.57<sup>a</sup> | 73.19<sup>a</sup> | 74.61<sup>a</sup> | 74.13<sup>a</sup> | 72.51<sup>a</sup> | 70.37<sup>b</sup> | 0.66 |
| Protein | 13.40 | 13.95 | 15.22 | 14.31 | 15.90 | 14.21 | 14.68 | 0.30 |
| Lipid | 11.00 | 8.76<sup>e</sup> | 9.62<sup>c</sup> | 8.48<sup>d</sup> | 8.7<sup>e</sup> | 11.07<sup>b</sup> | 13.62<sup>a</sup> | 0.81 |
| Ash | 3.30 | 2.68<sup>a</sup> | 2.09<sup>c</sup> | 2.24<sup>b</sup> | 2.27<sup>b</sup> | 2.12<sup>c</sup> | 2.19<sup>b</sup> | 0.09 |

A higher ash content in the tilapia fed on the control (CON) fish meal diet was obtained due to the high-level ash content (minerals) in the fish meal, as can be referred to in Table 2. Moreover, these fish also contained an elevated level of moisture content. Body protein and ash were unlikely to be influenced when the size of the fish was taken into consideration. After eight weeks of the trial period, all those groups of fish were observed to have an increase in their carcass protein level. However, fish fed on the fish meal based control diet CON, SESB50, SESB75 and FFSB50 diets showed a lower lipid level in their final body composition compared to the initial lipid content of tilapia.

## Discussion

The effect of alternative protein sources and the replacement of fish meal for commercially valuable farmed fish and a strategy for modern aquafeeds has been extensively studied (Maas *et al*., 2020; Gule and Geremew, 2022). It was concluded that increasing plant protein levels to replace fish meal may have detrimental effect on growth rate and feed utilisation at higher levels. In the present work, the results obtained demonstrate that plant protein sources can contribute up to 50% within a high-quality fish meal based diet for tilapia without much reduction in growth rate, particularly the soybean meal. This agrees with the results previously obtained by Sadiku and Jauncey (1995) and Gallagher *et al*. (1994). Daniel (2018) presented a review of the major plant ingredients that could be combined to make even a 100% replacement of fish meal in diets for fish, including tilapia, feasible. It was stated that plant ingredients generally have appreciable levels of Anti-Nutritional Factors (ANFs) that are adversely bioactive in fish. Plant protein concentrates are also deemed deficient in specific EAAs, such as methionine and lysine and with lower overall nutrient digestibility, resulting in inferior nutrient bioavailability (Daniel, 2018).

Higher substitution levels with plant proteins in this investigation markedly reduced the growth rate and performance of tilapia above 50% incorporation of both soybean meal and corn gluten meal at the expense of LT fish meal and wheat. The possible reason for the poor performance of high substitution of plant proteins in diets is the imbalance of their nutrients. The three factors which are likely to produce the decreased growth rates observed at higher protein inclusion levels are (1) toxic components such as trypsin inhibitor protein in soybean and digestive disturbances due to soya oligosaccharides (Rumsey, 1993). (2) lower digestibility of plant proteins and carbohydrates. (3) limiting of essential amino acids particularly methionine (in soybean) and lysine (maize gluten).

Of the various antinutritional factors (ANFs), great importance is attached to those which interfere with the digestion and absorption of protein. These inhibit the proteolytic activity of certain enzymes and are hence termed protease inhibitors. To minimise possible hazards and improve soybean meal nutritional quality, inhibitors could be generally inactivated by heat treatment or aqueous or solvent extraction processes. There have been studies previously to evaluate soybean in fish diets, as a practical ingredient. Earlier research by Webster Goodgame-Tiu and Tidwell, (1995) on blue catfish, reported that the use of heated SBM did not increase the growth of the fish, probably because of the already low level of trypsin inhibitor in the commercial SBM used. On the other hand, higher growth was obtained in Nile tilapia, fed diets containing heated treated (boiled) SBM compared with fish fed diets containing fish meal or raw SBM (Wee and Wang, 1987). This may be due to the inactivation of the high trypsin inhibitor activity in the SBM and the increasing digestibility of the diets containing boiled SBM compared to raw SBM.

Poor performance at the higher levels of plant protein is mostly likely due to the low methionine level. However, another factor should be considered, such as the incomplete denaturing of trypsin inhibitors and haemagglutinins during the processing treatment of the meal. More recently, Sharda, Sharma, and Saini (2017) investigated fish meal replacement with Nile tilapia (*Oreochromis niloticus*), indicating that an inclusion level of just 25% was feasible in experimental diets. However, Ahmad *et al*. (2020) was able to confirm that 50% replacement of fish meal was achievable in their studies with juvenile tilapia (*Oreochromis niloticus*) reared in saline conditions with a dietary protein level of 35%, similar to the current investigation.

In the present study, the percentage of dietary protein was set at 38% and Jauncey (2000) gave more precise information on the nutrient requirements and essential amino requirements for tilapia over a range of weight categories. No supplementary amino acids were included in the experimental formulations in this current study with tilapia. It is also noted that no effect on growth and food utilisation of *0. niloticus* fry when 75% of herring meal in the basal diet was replaced by FFSB supplemented with di-methionine (Shiau, Chuang and Sun, 1987). More recently, Pervin *et al*. (2020) found that up to 75% of fish meal in diets for tilapia could be replaced with soybean meal, validating our results. These earlier workers found that very high inclusions above this margin caused shortening of the gut villi length and other adverse histomorphology changes. Intestinal submuscularis thickness was found to be inversely related with the increasing villus height. Also, proteolytic activity significantly elevated the stomach, mid-intestine, and distal gut of fed with SBM0 compared with SBM100 dietary inclusion.

Similar to salmon, tilapia may be sensitive to certain types of soybean meal, causing enteritis-like conditions. Another possible explanation for the inferior growth, particularly at 75% inclusion of solvent-extracted soybean meal with the fish meal, is the low digestibility bioavailability of minerals, especially phosphorus, compared with fish meal (Nguyen, 2008). In the present work, our study did not include mineral bioavailability or retention measurements as this investigation focused on protein assimilation.

Fish have a high requirement for phosphorus. However, soybean is deficient in available phosphorus. Although SBM contains approximately 0.7% phosphorus (P), only about one-half is biologically available to fish (Lovell, 1988). However, this could be effectively corrected by dietary supplementation with inorganic phosphorus sources such as dicalcium phosphate. More recently, several workers have reported the effective use of phytase as an exogenous enzyme to increase P utilisation in fish, including tilapia (Nwanna and Schwarz, 2006; Jiang *et al*. 2014). Adeoye *et al*. (2016) examined a suite of mixed enzymes from a solid-state fermentation (SSF) commercial feed additive with positive results for plant inclusion in diets for tilapia. In the current experiment, high inclusion of soybean meal may have compromised P requirements of tilapia and mineral supplementation may not have been sufficient.

The present study primarily replaced fish meal with the plant ingredients (solvent-extracted soybean meal, full fat soya, and maize gluten meal) and wheat meal, accommodating varying carbohydrate contributions to the test diets. Studies reporting the digestibility of carbohydrates, starch, and especially non-starch polysaccharides (NSP) in tilapia are scarce. Carbohydrate digestibility in the diet is mainly associated with carbohydrate composition (starch vs. NSP). NSP (non-starch polysaccharides) are often considered to be indigestible and consequently of minor nutritional value. This has been reviewed by Maas *et al*. (2020), but it is stated by these authors that attention to using various feed enzymes, as mentioned, can lead to elevated digestibility and more available dietary energy. Digestibility of soybean meal and full fat soybean as well as maize gluten meal was reported by Davies *et al*. (2011) for tilapia and they determined high digestibility for energy, protein, and individual essential amino acids. These were over 83% on average for apparent digestibility for protein in each ingredient tested.

Palatability might also have hindered the growth rate in the higher replacement of fish meal with plant proteins. In this present study, reduced diet acceptability may be a reason for the significant difference in growth performance between the groups of experimental fish. This is clearly indicated by the significant difference in the daily food intake, especially with the higher inclusion of plant proteins. Some fish, such as red drum, find soybean meal unpalatable and will not consume diets without fish meal (Webster Goodgame-Tiu and Tidwell, 1995). Mohsen and Lovell (1990) reported that adding animal by-products to an SBM-based diet improved palatability for channel catfish. In the current trial, tilapia found the SESB75 diet to be unpalatable, as indicated by the poor feed efficiency value and low feed intake.

Fish fed on the MG50 diet did not differ significantly from the SESB75 diet regarding growth performance and nutrient utilisation. It has been reported that the performance of carp and tilapia decreased when fed pelleted diets containing 65-75% of maize grains. Al-Asgah and Ali (1994) reported that maize grain at 25% level in the tilapia diet showed better growth performance and energy utilisation than other carbohydrate sources like starch, dextrin, sucrose, and glucose. Furthermore, the research made by Al-Ogaily (1996) on including maize grains (25-43%) in the diets of *O. niloticus* decreased their growth performance. However, this whole grain product is not comparable with the present study using a high protein concentrate gluten. The decrease in weight gain and specific growth rate in fish with the increasing level of maize grain in the diet might have also been due to a reduction in their protein content and, consequently, the protein-to-energy (P/E) ratio (El-Sayed and Teshima, 1991). Lysine is the limiting amino acid in maize gluten (De-Silva, 1997), which might be a possible explanation for the inferior growth performance obtained. It was also reported that arginine may also be a limiting amino acid for this ingredient and maize gluten showed a low digestible energy (DE), which was only recorded as 39% in carp. Low palatability is also a further factor that must be considered for the higher inclusion of maize gluten meal concerning poor feed intake. Kaur and Saxena (2005) also reported that the optimal maize gluten for catla (*Catla catla*), rohu (*Labeo rohita*) and mrigal (*Cirrhina mrigala*) carp species was at a 5% inclusion. These authors concluded that reduced digestibility and inferior lysine and methionine levels were primarily responsible for reduced growth performance and feed utilisation in these species.

Sales (2009) examined the effect of fish meal replacement by soybean products on fish growth based on meta-analysis for several fish species of commercial importance, which compares the digestibility of selected plant proteins, for example, solvent-extracted soybean, full fat soybean and maize gluten. These conclusions agree with the Apparent Net Protein Utilisation (ANPU) result in the current tilapia fed on SESB50 and FFSB50 diets. However, the inclusion of 50% solvent-extracted soybean and full fat soybean in the diet, still showed significantly reduced performance compared to the control diet. Recently, Iqbal, Yaqub and Ayub, (2022) reported a study to assess the effects of partial or full replacement of fish meal (FM) and soybean meal (SBM) with canola meal (CM) on growth performance, health status, and cost-benefit ratio of genetically improved farmed tilapia, *Oreochromis niloticus*. The findings of these workers agreed to our studies with similar size tilapia.

Protein Efficiency Ratio (PER) is the weight gain (biomass) ratio to dietary protein intake. This measures how effectively the fish utilise protein, but the expression includes the increase in biomass as a whole relative to dietary protein intake. The protein efficiency ratio of the control group (CON) tilapia was 2.68 compared to the fish fed on MG75, which was 1.34. This indicated a progressive reduction in nutrient assimilation, as can be observed from the lipid content in the carcass composition (Table 5). Furthermore, the trend shown by Apparent Net Protein Utilisation (ANPU) confirmed that elevated levels of plant proteins reduced protein retention efficiency as a direct measurement of protein utilisation and carcass deposition over the 8 weeks. High substitution of plant protein sources caused the lowest Net Energy Retention [Utilisation] (NEU) value. Additionally,the NEU results agreed with the poor feed intake by fish-fed on SESB75 and MG75 diets.

Knowledge of the body composition of fish, allows the assessment of fish health and determination of the efficiency of transfer of nutrients from the feed to the fish making it possible to predictably modify carcass composition (Al-Ogaily, 1996). The body composition results of tilapia in this experiment follow the growth performance expectations for this species. Tilapia fed the MG75 diet showed significantly lower (P < 0.05) body moisture, but higher fat levels. However, the lipid content in the final carcass was somewhat lower than the initial carcass in some fish groups. During the fry stage, fish increase in size rather than store energy when energy intake is limited. This generally leads to lower levels of lipids and higher levels of moisture. Whole body moisture is inversely related to whole body lipid and decreases or increases as lipid is stored or utilised (Table 5). In this study, growth and lipid content were affected by the diet. This study compared plant proteins only against a Low Temperature (LT) dried fish meal to serve the specific purpose of comparison to a protein of known high Biological Value (BV) with no nutritional constraints based on a control (clean diet) approach. Results of this study indicate that solvent-extracted soybean and full fat soybean meals as well as maize gluten meal are optimally utilized by tilapia fry at 50% contribution of the fish-meal-based diet. Higher levels (75%) produced inferior performance.

Further work is required to ascertain the use of other plant-based ingredients in more complex diet formulations for tilapia. The cost benefit of incorporating solvent-extracted soybean meal and full fat soybean meal as well as maize gluten meal needs to be factored for effective feed formulations.

## Acknowledgements

The authors thank Ms W. Tamat for her contribution to the practical aspects of this project and technical support.

## Ethical approval

The animal husbandry techniques involved in the study were undertaken under the supervision of Professor Simon Davies in accordance with the Animal (Scientific Procedures) Act of 1986, License #Pil: 30/00080. This was during Professor Davies’ previous tenure at The University of Plymouth.

